# Dysconnection of right parietal and frontal cortex in neglect syndrome

**DOI:** 10.1101/192583

**Authors:** Martin J. Dietz, Jørgen F. Nielsen, Andreas Roepstorff, Marta I. Garrido

## Abstract

A lesion to the right hemisphere of the brain often leads to perceptual neglect of the left side of the sensorium. The fact that lesions to different cortical regions lead to the same symptoms points to neglect as a dysconnection syndrome that may result from the dysconnection of a distributed network, rather than a disruption of computation in any particular brain region. To test this hypothesis, we used Bayesian analysis of effective connectivity based on electroencephalographic recordings in patients with left-sided neglect after a right-hemisphere lesion. While age-matched healthy controls showed a contralateral increase in connection strength between parietal and frontal cortex with respect to the laterality of the stimuli, neglect patients showed a dysconnection between parietal and frontal cortex in the right hemisphere when stimuli appeared on their neglected side, but preserved connectivity in the left hemisphere when stimuli appeared on their right. Crucially, this parieto-frontal feedback connectivity was aggravated in patients with more severe symptoms. In contrast, patients and controls did not show differences in the local connectivity within regions. These findings suggest that the aetiology of neglect may lie in the dysconnection of a distributed network, rather than the disruption of any particular brain region.

## Introduction

Neglect is one of the most common neurological disorders following stroke (Azouvi et al., 2002). Neglect patients are characterized by the fact that they ignore the side of the sensorium contralateral to a cortical or thalamo-reticular lesion, usually in the absence of lesions to primary sensory or motor areas (Molenberghs et al., 2012). Importantly, neglect of the left side of the sensorium after a right-hemisphere lesion is more frequent and severe than neglect of the right hemispace after a left-hemisphere lesion. This hemispheric asymmetry has been explained in terms of a right-hemisphere dominance of the fronto-parietal network that mediates attentional orienting to salient stimuli (Dietz et al., 2014). According to this hypothesis, a left-hemisphere circuit mediates attentional orienting to the right hemispace, whereas the right hemisphere is involved in the orienting to both sides of the sensorium (Mesulam, 1999). This is based on the existence of a dorsal fronto-parietal system for goal-directed deployment of attention and a ventral system for reorienting to salient stimuli that appear unexpectedly in the sensorium (Vossel et al., 2014). There is a general heterogeneity in the lesion profiles of neglect patients (Molenberghs et al., 2012) and the observation that a lesion to temporal, frontal or parietal cortex leads to a similar perceptual deficit points to the notion of neglect as a dysconnection syndrome (Lunven et al., 2015). While the dorsal system consists of the first and second branches of the superior longitudinal fasciculus, the ventral system connects the inferior frontal cortex with the inferior parietal lobule via the third branch of this fronto-parietal fascicle (de Schotten et al., 2011). Anatomical and behavioral evidence for a hemispheric asymmetry has been shown in healthy participants where the white matter volume of the third branch correlated with detection of targets in the left, but not the right visual field (de Schotten et al., 2011). Consistent with this relation between anatomy and behavior, neglect patients in the subacute phase show axonal degeneration in the second and third superior longitudinal fascicles of the right, but not the left hemisphere compared to stroke patients without neglect (Lunven et al., 2015). We recently provided evidence of a right-hemisphere dominance in the effective connetivity between the inferior frontal gyrus and the inferior parietal cortex in healthy individuals during unexpected left and right auditory stimuli (Dietz et al., 2014). However, whether patients with unilateral neglect have a right-hemisphere dysconnection has not yet been investigated. To this end, we recorded high-density electroencephalography (EEG) in ten patients with left-sided neglect following a right-hemisphere lesion and eleven healthy age-matched controls during an auditory oddball paradigm where the location of the stimulus changed unpredictably to the left or the right side in egocentric coordinates. Using dynamic causal modeling (DCM) of evoked electrophysiological responses (David et al., 2006), we analyzed the strength of feedforward and feedback connections between the primary auditory cortex in Heschl’s gyrus (HG), the inferior frontal gyrus (IFG) and the inferior parietal lobule (IPL). Within this hierarchy of cortical regions, we then tested a set of alternative hypotheses about the laterality of the ventral system that mediates perception of salient stimuli. Our set of hypotheses represented (a) a contralateral coding of the sensorium with respect to the side of the stimulus (b) a left-lateralized coding of the sensorium and (c) a right-hemisphere dominant coding (see Fig. 1). The hypothesis of a contralateral increase in effective connectivity was motivated by the “contralateral bias” model (Kinsbourne, 1987). The hypothesis that patients may show a left-lateralized cortical network was based on their lesion profile in the right hemisphere that causes perceptual neglect of the left side of the sensorium. Finally, the hypothesis that the right hemisphere codes both sides of the sensorium was based on the right-hemisphere dominance model (Dietz et al., 2014; Mesulam, 1999).

**Figure 1.**
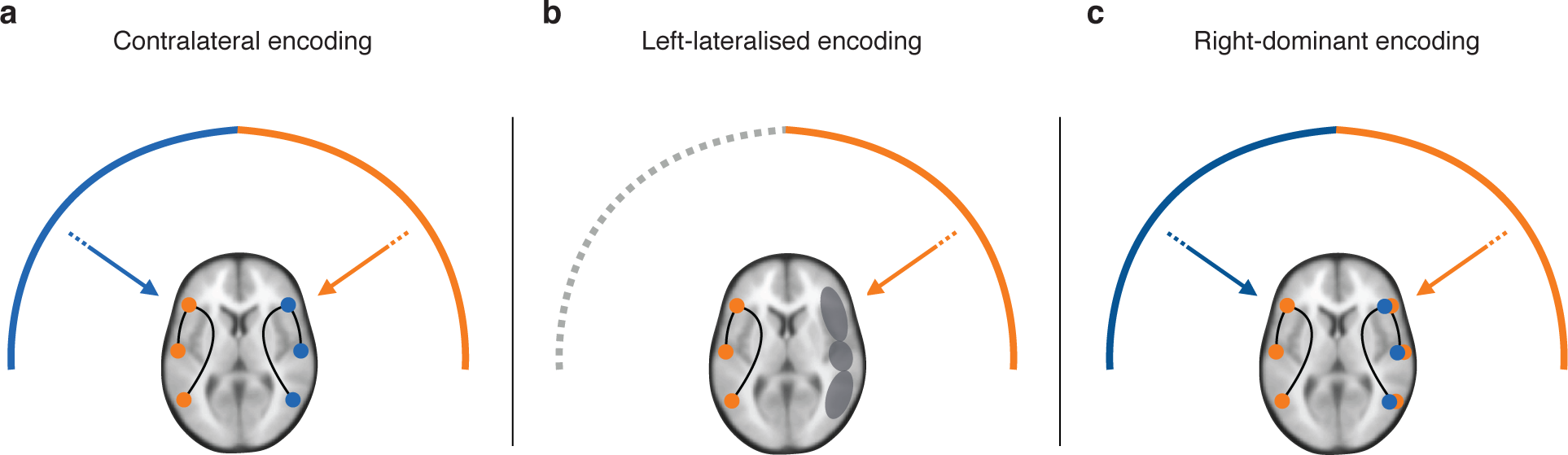
Alternative hypotheses about the coding of the sensorium in terms of the effective connectivity (synaptic efficacy) with respect to the side of the stimulus: (a) contralateral coding (b) left-lateralised coding and (c) right-dominant coding. Each network was analyzed as a hierarchy consisting of feedforward and feedback connections between Heschl’s gyrus (HG), the inferior frontal gyrus (IFG) and the inferior parietal lobule (IPL) in the relevant left or right hemisphere.

## Methods

### Experimental design

We recorded 64-channel electroencephalography (EEG) from patients and healthy controls using active Ag/AgCl electrodes placed according to the extended 10-20% system (Brain Products, GmbH). Data were sampled at 1 kHz with a 0.1 Hz high-pass filter. Participants were presented with an auditory oddball paradigm where the location of a tone changed in an unpredictable fashion from the midline to the left or the right side in egocentric coordinates. Stimuli perceived as originating from the midline were repeated consequently between 3-7 times by randomly sampling from a deterministic multinomial distribution, thus occurring with 80% probability. An interaural time delay (ITD) of 800 *μ*s between left and right ears was used to create a change in the location of the stimulus to the egocentric left and right side at approximately 90° angle in the horizontal plane. Stimuli at left and right locations each had 10% probability of occurrence. All other spectral, amplitude and duration parameters were kept constant. Stimuli consisted of sinusoidal pure tones with 75 ms duration, including 5 ms fade-in and 5 ms fade-out. We used Presentation software (Neurobehavioural Systems, Inc) to deliver stereo stimuli through in-ear headphones (Sennheiser, GmbH) at a stimulus-onset asynchrony of 500 ms. The sound pressure level was manually adjusted to lie between the individual participants’ hearing threshold and their nuisance level. Each participant was presented with a total of 1800 stimuli (1440 midline, 180 left and 180 right). Participants were instructed to maintain fixation on a central cross (on a black screen) during the experiment. Ethical approval was obtained from the local ethics committee of the Central Denmark Region, Denmark and patients and healthy controls gave their informed written consent before the experiment according to the Declaration of Helsinki.

### EEG data analysis

Data analysis was performed using Statistical Parametric Mapping (SPM12) academic freeware implemented in Matlab (MathWorks Inc., USA). The EEG data were re-referenced to the average over channels, down-sampled from 1 kHz to 250 Hz, high-pass filtered at 0.5 Hz and low-pass filtered at 30 Hz using a two-pass Butterworth filter. Experimental trials were epoched from −100 to 400 ms in peri-stimulus time and baseline-corrected using the average over the pre-stimulus time-window. Artefacts were rejected by thresholding the signal at 80 *μ*V, leaving on average 80% of trials for analysis. Trials were averaged in the time-domain using robust averaging to form evoked responses. To enable analysis in sensor space over post-stimulus time, the evoked dataset was converted into a 3-dimensional spatiotemporal image consisting of a 2D sensor representation over time. Images were smoothed with a Gaussian kernel of 20 mm full-width at half maximum (FWHM) in the spatial dimension and 20 ms in the temporal dimension. The results were thresholded at *p <* 0.05, corrected for the family-wise error (FWE) rate using random field theory (Kilner and Friston, 2010).

### Specification of dynamic causal models

We specified a set of dynamic causal models that describe the effective connectivity between temporal, frontal and parietal areas. These models tested a set of alternative hypotheses about the laterality of the cortical mechanisms that mediate the perception of a stimulus in a new spatial location. Our alternative hypotheses represented a contralateral increase in the strength of effective connectivity with respect to the side of the stimulus (model C), a left-lateralization in effective connectivity (model L) and a right-hemisphere dominance (model R). Following our previous study in normal subjects (Romanski et al., 1999), we used existing anatomical knowledge to set the prior mean of the dipole locations - used to summarize each cortical area - to the following coordinates in MNI standard space: left [−42 −22 7] and right [46 −14 8] Heschl’s gyrus (HG), left [−46 28 8] and right [46 28 8] inferior frontal gyrus (IFG), left [−49 −38 38] and right [57 −38 42] inferior parietal cortex (IPC). The cortical network received afferent feedforward input to primary auditory cortex via the medial geniculate nucleus of the thalamus. This was modelled as a Gaussian impulse with a prior mean of 60 ms and prior standard deviation of 16 ms. The primary auditory cortex connected to the inferior frontal gyrus, which in turn connected to the inferior parietal cortex via feedforward and feedback connections. We modelled the data during the post-stimulus period 0-350 ms. This period encompasses components of the evoked response that are assumed to reflect the detection of a change in the stimulus location (Dietz et al., 2014) and a subsequent bottom-up attentional reorienting to the left or the right hemispace.

### Dynamic causal modelling of evoked responses

Dynamic causal modelling (DCM) is a method for estimating the directed coupling between sources of a cortical network and how this coupling changes with stimulus context. This context-dependent connectivity is referred to as effective connectivity and defined as the influence one neuronal population exerts on another population at the synaptic level. This renders dynamic causal modelling a generative model of the mechanisms that generate observed responses, as opposed to metrics of functional connectivity that operate at the level of data features, such as coherence, phase-locking or Granger causality. We used a biologically realistic neural-mass model (David et al., 2006) that summarizes the dynamics, within an electromagnetic source, as the average depolarization of four cell populations whose intrinsic connections, within a cortical column, conform to a canonical cortical microcircuit (CMC) (Bastos et al., 2012). Each cortical source comprises a population of excitatory spiny stellate cells (1), inhibitory interneurons (2), and distinct populations of excitatory pyramidal cells in deep layers (3) and superficial layers (4) that are the sources of feedback and feedforward and feedback connections between sources, respectively (see Fig. 4a for a schematic of the CMC). This dynamic causal model of evoked electrophysiological responses is specified by a set of coupled differential equations that model the dynamics of depolarization *V* as a function of time

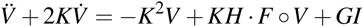

This model predicts how the average depolarization *V* = (*v*_1_*, v*_2_*, v*_3_*, v*_4_)^T^ of each neuronal population *v*_*i*_(*t*) evolves in time as a function of depolarisation in another population *v* _*j*_(*t*) and stimulus-evoked input *I*(*t*). The first term describes the decay of post-synaptic responses, the second term is a mapping from post-synaptic membrane depolarization to pre-synaptic firing rate and a convolution of pre-synaptic firing to cause post-synaptic depolarization in the target population. The last term describes the stimulus-evoked input to the system. The parameters *H* = *diag*(*h*_*e*_*, h*_*i*_*, h*_*e*_*, h*_*e*_) control the maximum post-synaptic response of the excitatory and inhibitory populations, modelled as an alpha-function kernel with time constants *K* = (*k*_1_*, k*_2_*, k*_3_*, k*_4_) that describe their rate of post-synaptic decay. *F* is the nonlinear mapping from post-synaptic depolarization *V* to pre-synaptic firing rate that serves as input to the target population. *G* = (*k*_1_*h*_*e*_, 0, 0, 0)^T^ maps from thalamic input *I* to the depolarisation of spiny stellate cells in the granular layer. The rate of change of each population in the microcircuit is then given by

Spiny stellate cells (1)

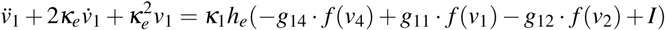

Inhibitory interneurons (2)

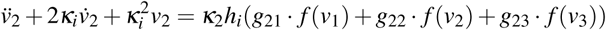

Deep pyramidal cells (3)

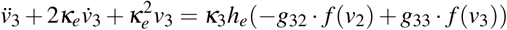

Superficial pyramidal cells (4)

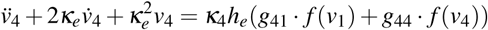

where *g*_*ij*_ is the connection strength from the *j*th to the *i*th population and *f* (*v* _*j*_) is the nonlinear sigmoid function

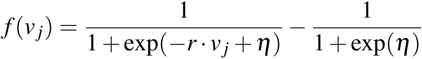

that transforms excitatory or inhibitory post-synaptic depolarization into an average density of pre-synaptic firing. The slope *r* of the sigmoid function controls the synaptic gain and *h* is the threshold for eliciting deviations from baseline firing. The model is parameterised with a log-scale parameter on its intrinsic and extrinsic feedforward and feedback connections to provide posterior estimates of the connection strengths *p*(*ϑ y, m*), given electrophysiological measurements *y*. An observation model then maps the predicted responses of microcircuit to observed channel data using a measurement-specific gain function *g*(*V, ϑ,φ,*) : *V → Y*. We used a spatiotemporal formulation of a conventional equivalent current dipole (ECD) forward model

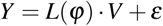

This models observed channel data *Y* as a mixture of post-synaptic depolarization *V* and additive Gaussian error *e*. Each source in the network was modelled as a dipole whose locations and orientations φ parameterize the electromagnetic lead field *L* that describes the spatial mapping from neuronal depolarisation *V* to observed channel data *Y*, assuming additive Gaussian measurement noise. This lead field was based on a boundary element method (BEM) head model to describe the propagation of the electrical current through the tissues and skull onto the scalp surface (Mosher et al., 1999). This forward model assumes normal volume conductance of the electrical current through the tissue and cranial layers when predicting the observed scalp topography. Openings in the skull have a distortive effect on the propagation of the electrical potential generated by post-synaptic currents (Wolters et al., 2006). This means that unmodelled violations of normal volume conductance would invalidate source reconstruction of scalp EEG data within patients and hence create spurious results between patients and healthy controls. For this reason, we only included patients who had not undergone craniotomy.

### Variational Bayesian estimation

Dynamic causal models are estimated using variational Bayes (Friston et al. 2007). Given a model *m*, specified in terms of the prior distribution *p*(*ϑ|m*) of the model parameters and a likelihood function *p*(*y |ϑ, m*) for observed data *y*, this provides both the posterior distribution of the connection strengths *p*(*ϑ|y, m*) and the marginal likelihood of the model itself, also known as the model evidence

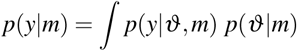

Using a Laplace approximation to the posterior distribution of the parameters *q*(*ϑ*) ∼*N* (*μ,* ∑), the conditional means and covariances of the parameters are estimated iteratively by maximizing a lower bound on the logarithm of the model evidence ln *p*(*y|m*). This optimisation is formulated as a Newton ascent on the (negative) free energy *F* of the model

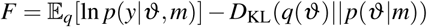

This renders the free energy an approximation to the log-evidence, which is exact when *q*(*ϑ*)= *p*(*ϑ|y, m*). The first term of the free energy is the expected log-likelihood of data y and represents the accuracy of the model. The second term is the relative entropy or Kullback-Leibner divergence between the multivariate posterior and prior probability densities and represents the complexity of the model. The formulation of the complexity as a divergence rests on the assumption that the posterior should not have to move too far from the prior to accommodate the data. In other words, a good model should provide posterior beliefs that are consistent with its prior beliefs after observing new data. The free energy can thus be decomposed into accuracy minus complexity. Together, these two terms summarize the quality of a model in explaining the data.

### Bayesian model selection

Dynamic causal models are compared using their model evidence (Penny 2012). We used a random-effects Bayesian model selection procedure to provide the posterior probability and protected exceedance probability of each model for model comparison within each population (Penny et al. 2010). The protected exceedance probability was recently introduced in the statistical literature in the context of Bayesian model selection to quantify the posterior probability that a model is more frequent in the sample than the other models in the comparison set, above and beyond differences in model frequencies that are expected to occur by chance (Rigoux et al. 2014).

## Results

### Patient data

We recorded EEG responses and behavioural data from 10 patients (mean age 59, age range 41-68) with neglect of the left side of the sensorium during their subacute phase. The control group was age-matched to the patient group to within a few years and consisted of 11 healthy volunteers (mean age 56, age range 44-71) with no history of neurological or psychiatric disorder. The patients had heterogeneous lesion profiles in the right cerebral hemisphere and the severity of contralesional neglect was classified according to the Behavioural Inattention Test (BIT) (Wilson et al., 1987). The BIT scale ranges from 0-227, where a lower score is taken as evidence of more severe neglect. All patients expressed neglect symptoms corresponding to a level of severity below the conventional diagnostic criterion of 129 points at the time of EEG acquisition. To ensure the validity of the electrophysiological data, we only included patients who had not undergone craniotomy. This is because (unmodelled) violations of normal volume conductance will invalidate source reconstruction of EEG data in patients and hence create spurious results between patients and healthy controls.

### Evoked responses in healthy participants

We first tested for cortical responses evoked by salient left and right stimuli in healthy participants using one-sample t-tests at the group level, family-wise error (FWE) corrected for multiple comparisons using random field theory (Kilner and Friston, 2010). This analysis revealed evoked responses over the fronto-central scalp to both left and right stimuli, relative to stimuli presented in the midline (Fig. 2a) which resemble the typical mismatch negativity (MMN) observed in oddball paradigms. These responses peaked at 164 ms for the left (*t*(10) = 9.89, *p* = 0.003, FWE-corrected one-sample *t*-test) and at 176 ms for the right stimulus (*t*(10) = 7.39, *p* = 0.005, FWE-corrected one-sample *t*-test). Later in post-stimulus time, we observed a response over the parieto-central scalp, evoked by left and right stimuli with a peak at 260 ms for the left (*t*(10) = 4.5, *p* = 0.001, FWE-corrected one-sample *t*-test, Fig. 2b) and a peak at 276 ms for the right stimulus (*t*(10) = 2.89, *p*= 0.009, FWE-corrected one-sample *t*-test, Fig. 2c). We interpret this positivity as a P3a, which is typical in probabilistic oddball paradigms that induce reorienting of attention to salient stimuli (Polich, 2007).

**Figure 2.**
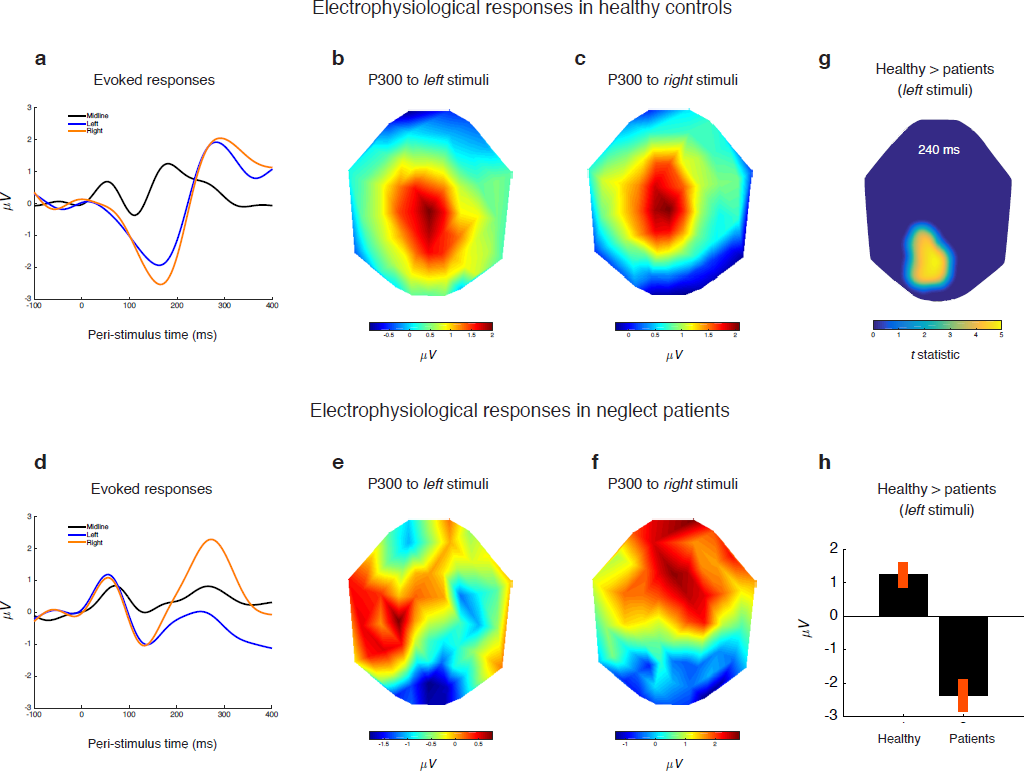
Evoked cortical responses in healthy participants and neglect patients (a) Evoked responses to midline, left and right stimuli in healthy participants (b) Scalp topography of the P3a induced by left stimuli at 260 ms post-stimulus time (c) Scalp topography of the P3a induced by right stimuli at 276 ms post-stimulus time (d) Evoked responses to midline, left and right stimuli in neglect patients (e) Scalp topography induced by stimuli on the patients’ neglected side at 292 ms post-stimulus time (f) Scalp topography of the P3a induced by right stimuli at 292 ms post-stimulus time (g) Statistical t-map of the difference between healthy participants and neglect patients during left stimuli with a peak at 240 ms post-stimulus time (h) Average responses and 95% confidence intervals in healthy participants and neglect patients at the peak of the parieto-central cluster identified in (g) above.

### Evoked responses in neglect patients

Having established a typical MMN and P3a in healthy controls, we then tested for cortical responses evoked by salient left and right stimuli in neglect patients using a one-sample *t*-test at the group level, corrected for multiple comparisons. This revealed evoked responses to both left and right stimuli, relative to midline stimuli (Fig. 2d) that again resembled a MMN with a peak at 128 ms for the left (*t*(9) = 4.87, *p* = 0.007, FWE-corrected one-sample *t*-test) and a peak at 156 ms for the right stimulus (*t*(9) = 5.72, *p* = 0.04, FWE-corrected one-sample *t*-test). We then tested for responses later in post-stimulus time and found an evoked response when salient stimuli appeared on the patients’ right side, with a peak at 292 ms (*t*(9) = 2.25, *p* = 0.02, FWE-corrected one-sample *t*-test, Fig. 2e). Again, we interpret this positivity as a P3a, which is known to reflect attentional reorienting to salient or unexpected stimuli in healthy participants (Polich, 2007). However, when stimuli appeared on the left (neglected) side, there was no significant response at this latency (Fig. 2f). To confirm the absence of a P3a in neglect patients compared to the P3a observed in healthy controls, we tested for a difference in evoked responses between controls and patients and found a significant difference over the parieto-central scalp with a peak at 240 ms (*t*(19) = 4.88, *p* = 0.02, FWE-corrected two-sample *t*-test, Fig. 2g and 2h).

### Bayesian analysis of effective connectivity

We then analyzed the electrophysiological responses using dynamic causal modeling (DCM). Following our previous study in neurotypical participants (Dietz et al., 2014), we used existing anatomical knowledge to set the prior mean of the dipole locations, used to summarize each cortical region, to the following coordinates in MNI space: left [−42 −22 7] and right [46 −14 8] Heschl’s gyrus (HG), left [−46 28 8] and right [46 28 8] inferior frontal gyrus (IFG), left [−49 −38 38] and right [57 −38 42] inferior parietal cortex (IPC). This cortical network received feedforward input, modelled as a Gaussian impulse, to primary auditory cortex via the medial geniculate nucleus of the thalamus (Kaas and Hackett, 2000). The primary areas on the superior temporal plane connected to the inferior frontal gyrus (Romanski et al., 1999) which connected to the inferior parietal cortex via feedforward and feedback connections (de Schotten et al., 2011). This analysis provided the posterior density of the strength of feedforward and feedback connections in each patient and control between HG, IFG and in each hemisphere during salient left and right stimuli, relative to the midline. In addition to these extrinsic connections between sources, we also analyzed the strength of intrinsic connectivity within each cortical source. The posterior distribution of the connection strengths are estimated by maximizing a lower bound on the log-likelihood of the network model itself using variational Bayes (Friston et al., 2007). This provides the Laplace free energy of each model as an approximation to its log-likelihood, also known as the model evidence. Using Bayesian model selection, we then compared the log-evidences for our alternative hypotheses: (a) contralateral coding (b) left-lateralised coding and (c) right-dominant coding in patients and controls, separately. This allowed us to identify distinct network architectures for patients and controls under the assumption that patients use a reduced cortical network due to their right-hemisphere lesion.

### Right-dominant network in the typical brain

Consistent with our previous results in healthy younger adults (aged 20-35 years) (Dietz et al., 2014), the healthy controls (aged 44-71 years) showed a right-hemisphere dominance in the effective connectivity between temporal, frontal and parietal cortex (Protected exceedance probability *P >* 0.88, Fig. 3b). We then tested for a significant increase in connection strengths within this hierarchical network during left and right salient stimuli. An increase in connection strength corresponds to an increase in post-synaptic efficacy or the influence of an axonal projection on its post-synaptic target population. This analysis revealed that, when salient stimuli appeared on the left, there was an increase of 47% in the strength of the feedforward connection from right IFG to IPL (*t*(10) = 3.3, *p* = 0.004, one-sample *t*-test, Bonferroni-corrected). This feedforward influence was reciprocated by an increase of 28% in the strength of the feedback connection from right IPL to IFG (*t*(10) = 2.9, *p* = 0.007, one-sample *t*-test, Bonferroni-corrected) and an increase of 91% in the feedback connection from right IFG to HG (*t*(10) = 2.9, *p* = 0.007, one-sample *t*-test, Bonferroni-corrected). In contrast, when salient stimuli appeared on the right, there was an increase of 133% in the strength of the feedforward connection from left IFG to IPL (*t*(10) = 3.3, *p* = 0.004, one-sample *t*-test, Bonferroni-corrected). Similarly, this increase in the feedforward influence was reciprocated by an increase of 76% in the strength of the feedback connection from left IPL to IFG (*t*(10) = 2.2, *p* = 0.02, one-sample *t*-test) and an increase of 83% in the feedback connection from left IFG to HG (*t*(10) = 2.1, *p* = 0.03, one-sample *t*-test). Finally, there was an increase of 74% in the strength of the feedback connection from IFG to HG in the right hemisphere (*t*(10) = 1.9, *p* = 0.03, one-sample *t*-test) (see Fig. 3d). In other words, when stimuli appeared on the left or the right side in egocentric coordinates, there was an increase in the strength of feedforward and feedback connectivity in the hemisphere contralateral to the side of stimulation, with an additional feedback influence in the right hemisphere when stimuli appeared on the right. This replicates our earlier findings in the younger adult brain that showed a similar contralateral coding of stimulus laterality within a right-dominant network (Dietz et al., 2014). Finally, as a complement to Bayesian model comparison, we assessed the accuracy of the optimal model in explaining the observed data in terms of the difference between the model’s predicted responses and the observed electrophysiological responses. This showed that the right-dominant DCM explained, on average, 59% of the variance in the healthy controls.

**Figure 3.**
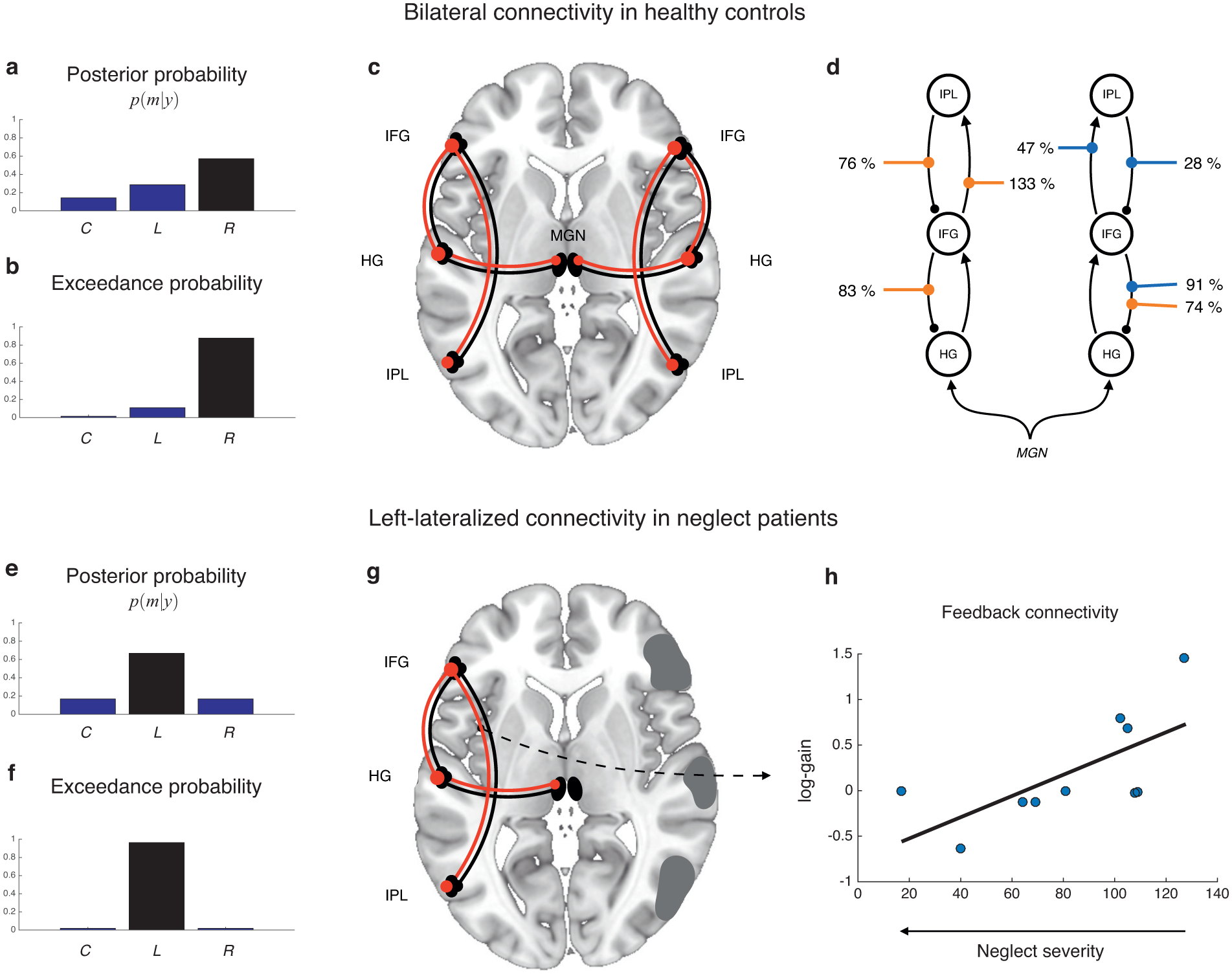
Right-dominant network in healthy controls and left-lateralised network in neglect patients (a) The posterior probability of the alternative network models representing contralateral coding (C), left-lateralised coding (L) and right-dominant coding (R) in healthy controls (b) Protected exceedance probability of the alternative network models in healthy controls (c) Architecture of the right-dominant cortical hierarchy with feedforward connections in black and feedback connections in red (d) Network graph showing contralateral increases in connection strength (%) during left stimuli indicated in blue and during right stimuli indicated in orange (e) The posterior probability of the alternative network models in neglect patients (f) Protected exceedance probability of the alternative network models in neglect patients (g) Architecture of the left-lateralised cortical network with feedforward connections in black and feedback connections in red (d) Linear relation between the severity of neglect symptoms (BIT score) between patients and their connection strengths (log-gain) during stimuli on the neglected side of the sensorium.

### Left-lateralized network in neglect patients

We then analyzed the evoked electrophysiological responses in neglect patients to identify abnormalities in their cortical mechanisms that may explain the reduced P3a response when stimuli appeared on their left (neglected) side, compared to their right (intact) side of the sensorium. This revealed a left-lateralized change in the effective connectivity between temporal, frontal and parietal cortex during both left and right salient stimuli (Protected exceedance probability *P >* 0.95). The reliability of this result is supported a Bayesian omnibus risk of *P <* 0.0001 which represents the probability that the observed sample of patients who use a left-lateralised network occurred by chance (Rigoux et al., 2014). This left-lateralization of the effective connectivity is entirely consistent with the patients’ lesion profiles that were all restricted to the right hemisphere. Although the lesions were anatomically heterogeneous among patients and included lesions in temporal, frontal and parietal cortex, the left-lateralization of effective connectivity was remarkably consistent across patients, hence providing evidence for neglect as a dysconnection syndrome. We then tested for significant changes in connection strengths within this left-hemispheric network. This revealed that, when salient stimuli appeared on the patients’ right side, there was an increase of 86% in the strength of the feedback connection from left IPL to IFG (*t*(9) = 2.4, *p* = 0.01, one-sample *t*-test, Bonferroni-corrected). This mirrors the increase in the strength of the corresponding homotopic connection in the healthy controls and is consistent with the patients’ preserved perception of the right side of the sensorium. In contrast, when the stimuli appeared on their left side, there was an increase of 120% in the strength of the feedback connection from left IFG to HG (*t*(9) = 1.96, *p* = 0.04, one-sample *t*-test), indicating perhaps a compensatory mechanism for ipsilateral coding in this sample of patients.

### Patients with more severe symptoms have weaker feedback connectivity

We then tested for a relationship between the patients’ individual differences in neglect severity, as expressed by their BIT score (Table 1), and the strength of cortical connections in response to stimuli on the patients’ left side. This revealed that patients with more severe symptoms had weaker feedback connectivity from the left IPL to IFG (*t*(8) = 3.09, *p* = 0.007, linear regression, Bonferroni-corrected). While caution is required when interpreting linear regression with a relatively small sample size, this finding suggests that neglect severity is associated with reduced top-down parieto-frontal connectivity. Note that stronger neglect symptoms correspond to a lower BIT score (Fig. 3h). In contrast, there was no evidence of such a relation between neglect severity and the strength of intrinsic connections within temporal, frontal or parietal areas. Again, we assessed the accuracy of the optimal model in explaining the observed data in terms of the difference between the model’s predicted responses and the observed electrophysiological responses. This showed that the left-lateralized DCM explained, on average, 64% of the variance in the patients’ data.

**Table 1.** ICH = intracranial haemorrhage, SAH = subarachnoidal haemorrhage, TBI = traumatic brain injury FIM = Functional Independence Measure (18-126), BIT = Behavioural Inattention Test (0-227)

| Patient | diagnosis | lesion | gender | age | months since event | FIM | BIT |
| --- | --- | --- | --- | --- | --- | --- | --- |
| 1 | ICH | parieto-occipital | male | 63 | 2 | - | 127 |
| 2 | SAH | temporal | male | 61 | 1 | 63 | 102 |
| 3 | infarction | temporo-parietal | male | 58 | 3 | 62 | 40 |
| 4 | infarction | temporo-occipital | male | 57 | 1 | 64 | 105 |
| 5 | SAH | temporo-occipital | female | 41 | 2 | 62 | 64 |
| 6 | SAH | temporo-occipital | female | 64 | 4 | 35 | 69 |
| 7 | ICH | parieto-occipital | male | 66 | 5 | 37 | 109 |
| 8 | SAH | temporal | female | 55 | 2 | 50 | 81 |
| 9 | TBI | frontal | male | 68 | 3 | 58 | 17 |
| 10 | infarction | fronto-parieto-temporal | female | 65 | 2 | 84 | 108 |

### Dysconnection of right parietal and frontal cortex in neglect patients

Having identified a left-lateralized network in neglect patients and a right-dominant network in the healthy subjects, we then tested for a difference in both extrinsic (cortico-cortical) and intrinsic (intra-cortical) connection strengths between the patients and the controls in both hemispheres. This was done using the Bayesian model average (BMA) obtained by weighting the average of each connection strength, within each subject, in relation to the posterior probability of each alternative model (Hoeting et al., 1999; Penny et al., 2010). This BMA effectively removes uncertainty about model structure and allows us to compare connection strengths between patients and healthy controls that use different networks in the brain. This revealed that when stimuli appeared on their left side, patients had weaker feedback connectivity from right IPL to IFG (*t*(19) = 2.04, *p* = 0.02, two-sample *t*-test), weaker feedforward connectivity from right IFG to IPL (*t*(19) = 1.89, *p* = 0.03, two-sample *t*-test) and weaker feedback connectivity from right IFG to HG (*t*(19) = 1.97, *p* = 0.03, two-sample *t*-test) compared to the age-matched healthy controls (see Fig. 4). This finding is a sign of impaired recurrent processing in the right hemisphere of neglect patients compared to healthy controls when stimuli appeared on their left side. In contrast, there was no evidence of a difference in connection strengths between patients and controls when stimuli appeared on their right side (significance criterion P ¡ 0.05, two-sample t-test). Finally, this also revealed that there is no evidence of a difference in the intrinsic connectivity between patients and controls (significance criterion *p <* 0.05, two-sample *t*-test, Fig. 4c).

**Figure 4.**
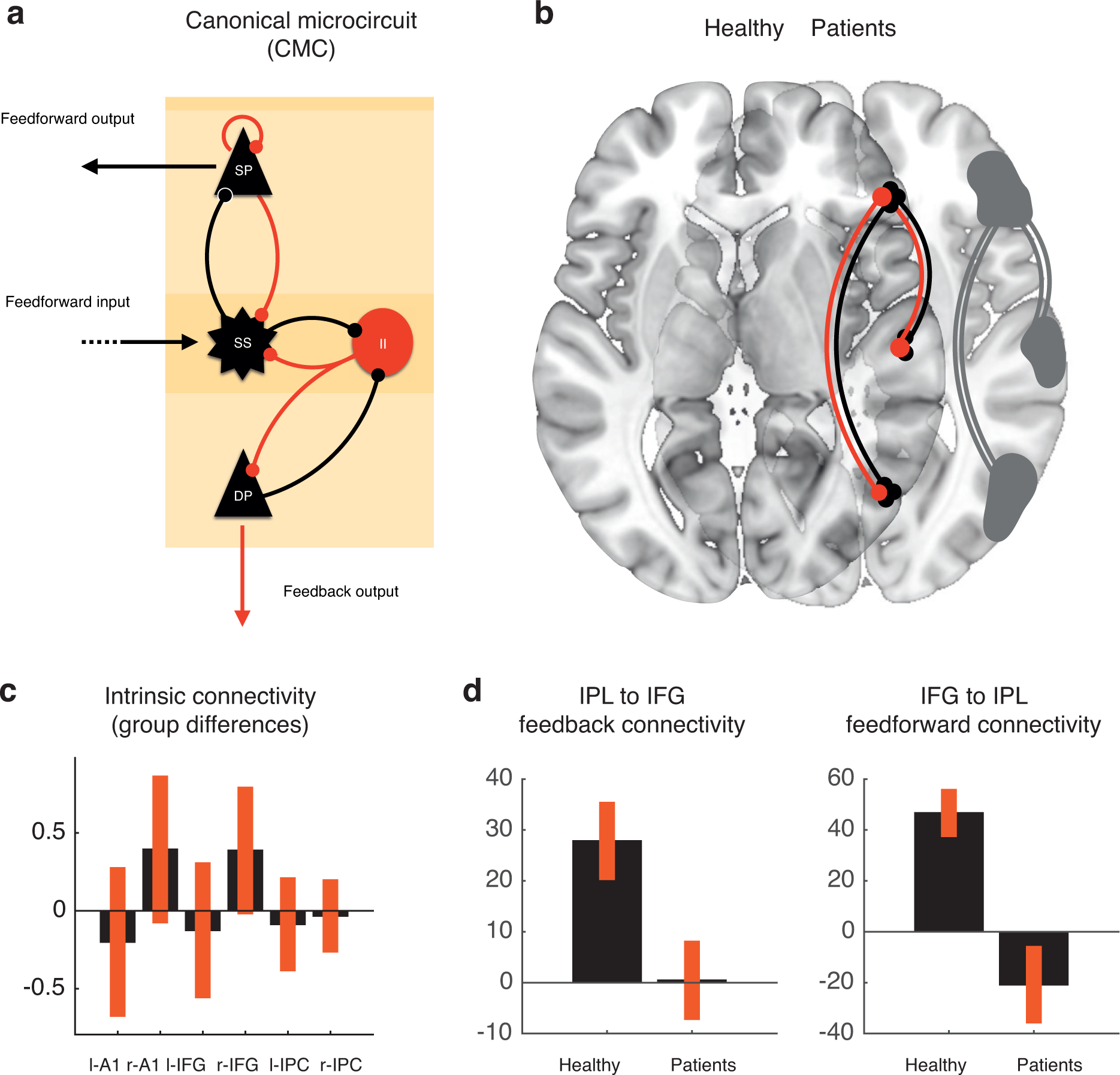
Dysconnection in neglect patients between parietal and frontal cortex in the right hemisphere during stimuli on the left side of the sensorium (a) Canonical cortical microcircuit (CMC) model used estimate the intrinsic and extrinsic connectivity within and between cortical sources. The CMC is a biologically realistic neural mass model comprising a population of spiny stellate (SS) cells in layer 4, a layer-wide population of inhibitory interneurons (II) and distinct populations of deep pyramidal (DP) neurons in layers 5/6 and superficial pyramidal (SP) cells in layers 2/3 that interact to generate feedforward and feedback output. (b) Fronto-parietal network in the right hemisphere that showed a difference in connection strengths between healthy controls and patients control. (c) Difference in connection strengths between patients and controls and 95% confidence intervals for the intrinsic connections within temporal, frontal and parietal areas of the network (d) Average connection strength and 95% confidence intervals for the feedback influence from right IPL to IFG and the corresponding feedforward connection, showing significantly weaker feedback influence in patients compared to healthy controls.

## Discussion

We used Bayesian analysis of effective connectivity to test a defined set of alternative hypotheses about the cortical network architecture that encodes the laterality of stimuli. This revealed that healthy controls use a right-dominant network. Within this bilateral network, we identified a contralateral increase in connection strengths with respect to stimulus laterality between hierarchically organized parietal, frontal and temporal regions. This finding replicates our previous work in younger adult brains using the same experimental paradigm (Dietz et al., 2014). Crucially, we show that neglect patients use a left-lateralized network with a dysconnection between parietal, frontal and temporal regions in the right hemisphere, which is aggravated in patients with more severe neglect. This is entirely consistent with their lesion profile, restricted to cortical lesions in the right hemisphere. Within this left-lateralized network, patients showed preserved feedback connectivity from parietal to frontal cortex when stimuli appeared on their right side. This mirrors the corresponding homotopic connection in the healthy controls and is consistent with the patients’ preserved perception of the right side of the sensorium. The presence of this feedback connection is also consistent with the presence of a P3a evoked response when stimuli appeared on their right side. In contrast, when stimuli appeared on their left side, a P3a response was absent compared to the healthy controls. This fits with the finding that neglect patients have a reduced P300 response to target stimuli in the neglected left visual field (Saevarsson et al., 2012). Using dynamic causal modelling, we were able to show that the responses elicited by auditory stimulation on the left side are explained by a dysconnection specific to the lesioned right hemisphere. Moreover, the absence of a P3a during left stimuli may be a consequence of this dysconnectivity between the inferior frontal gyrus and the inferior parietal lobule in the right hemisphere. This is consistent with the existing theories that the P3a is generated in the frontal and parietal cortex (Polich, 2007). Indeed, both patients with prefrontal lesions and patients with a lesion in the posterior parietal cortex show a reduction in the P3a to novel stimuli (Daffner et al., 2003).

### Dysconnectivity and diaschisis

Dysconnectivity may result from anatomical lesions to white matter bundles or as a consequence of a grey matter lesion that alters the synaptic efficacy of a population of neurons within its network. This is known as diaschisis and is classically associated with changes in neuronal excitability and decreased metabolism (Carrera and Tononi, 2014). More recently however, diaschisis has also been identified in the form of abnormal effective connectivity between regions due to a distant lesion (Boly et al., 2011; Campo et al., 2013). A lesion to an early area can thus affect a hierarchically higher area when feedforward afferents cease to innervate it. Similarly, a lesion in a higher area may affect a hierarchically lower area when feedback connections can no longer provide its simpler receptive fields with contextual modulation (Shipp, 2007). This is consistent with our finding that subacute neglect patients have weaker feedback connectivity from right IPL to IFG than healthy controls. Our findings thus show a systematic relation between decreased effective connectivity and perceptual symptoms, indicating that the feedforward and feedback connections that integrate parietal and frontal cortex may effectively encode information used by the brain to represent the location of stimuli in the sensorium.

### Neglect as a failure of predictive coding

It is conceivable that the organization of the brain as a hierarchy of columns and nuclei is dynamic and occurs whenever a population of neurons with a complex receptive field can predict the sensory features encoded by a population with a simpler receptive field, providing it with contextual information that can amplify or suppress the simpler features that it encodes. This is known as an extra-classical receptive field effect (Rao and Ballard, 1999) and has been shown in the macaque to depend on feedback connections (Hupé et al., 1998). In predictive coding, feedback connections are thought to provide lower-level populations of neurons with predictions in the form of prior beliefs about the sensorium, whereas feedforward connections mediate the ensuing prediction errors that result from the discrepancy between the brain’s predictions and changes in sensory input. The role of prediction errors is to update the representations of higher-level populations in order to provide Bayes-optimal predictions about the world. From a computational perspective, a dysconnection between areas that are hierarchically organized in this way will result in a Bayes-optimal loss of confidence in higher-level predictions about the sensorium. In this instance, higher-level populations can no longer predict the context of simpler stimulus representations encoded at lower levels. In other words, there is a failure of contextual modulation of simpler stimulus features via feedback mechanisms. In the presence of a new stimulus, the posterior parietal cortex may assume a hierarchically higher position in relation to the inferior frontal cortex (Dietz et al., 2014). In this stimulus context, the posterior parietal cortex may possess complex receptive fields that predict where stimuli can appear in the sensorium, encoded in egocentric coordinates (Andersen, 1997; Colby and Goldberg, 2003). When a stimulus occurs in a new location, like in our experiment, prediction errors are passed from the inferior frontal cortex to update the predictions about stimulus location encoded in the posterior parietal cortex. These distinct computational roles of feedforward and feedback connections may be reflected in their distinct anatomy and physiology: feedforward connections typically originate from pyramidal neurons in superficial layers 2/3 and target spiny stellate neurons in layer 4. In contrast, feedback connections typically originate in deep layers 5/6 and target diffusely in superficial and deep layers (Salin and Bullier, 1995; Shipp, 2007). These distinct, yet complementary roles of feedforward and feedback connections in perception are consistent with the weaker feedforward and feedback connectivity observed between parietal and frontal cortex in neglect patients. Our findings thus support the theory that both feedforward and feedback connections are necessary for perceptual inference and that both types of connections are impaired ipsilesionally in patients with unilateral neglect. Furthermore, the hierarchical organization that they confer on the brain may implement the hierarchical nature of predictive coding, where populations of neurons at a higher level are thought to encode predictions for sensory representations at lower levels of abstraction, encoded by populations with simpler receptive fields. These populations return their prediction errors in a feedforward fashion to update sensory perception at increasingly higher levels of abstraction. Indeed, this is in agreement with behavioural evidence of an general impairment of representational updating in neglect syndrome (Danckert et al., 2012).

In sum, we show that patients with a cortical lesion in the right hemisphere have a dysconnection between parietal and frontal cortex when stimuli appear on the neglected side of the sensorium, but preserved connectivity during stimuli on their right. At the same time, we show that the parietal to frontal feedback connectivity in the non-lesioned left hemisphere is aggravated in patients with more severe symptoms. In contrast, our analysis showed no difference in the intrinsic (intra-cortical) connectivity between patients and controls. This points to neglect as a dysconnection syndrome whose aetiology lies in the dysconnection of a distributed network, rather than the disruption of any particular brain region.

## Author contributions

M.J.D., M.I.G. and A.R. designed the experiment. J.F.N. provided the clinical and diagnostic information. M.J.D. performed the experiments in patients and healthy subjects. M.J.D. analyzed the data. M.J.D. and M.I.G. wrote the paper.

### Acknowledgements

This work was supported by the Danish Agency for Science, Technology and Innovation’s Investment Grant to MINDlab, Aarhus University. M.J.D is supported by the VELUX Foundation (00013930). M.I.G. is supported by a University of Queensland Fellowship (2016000071). We would like to thank neuropsychologists Mille Møller Thastum and Troels Jensen for their help with patient recruitment and neuropsychological testing.

